# TreeCluster: Massively scalable transmission clustering using phylogenetic trees

**DOI:** 10.1101/261354

**Authors:** Niema Moshiri

**Affiliations:** Bioinformatics and Systems Biology Graduate Program, UC San Diego, 9500 Gilman Dr., La Jolla, CA 92093,USA

**Keywords:** molecular epidemiology, network, transmission cluster, surveillance

## Abstract

**Background:** The ability to infer transmission clusters from molecular data is critical to designing and evaluating viral control strategies. Viral sequencing datasets are growing rapidly, but standard methods of transmission cluster inference do not scale well beyond thousands of sequences.

**Results:** I present TreeCluster, a cross-platform tool that performs transmission cluster inference on a given phylogenetic tree orders of magnitude faster than existing inference methods and supports multiple clustering optimization functions.

**Conclusions:** TreeCluster is a freely-available cross-platform open source Python 3 tool for inferring transmission clusters from phylogenetic trees. Code, usage information, and in-depth descriptions of the implemented clustering modes are available publicly at the following repository:

https://github.com/niemasd/TreeCluster

## Background

Epidemiologists often rely on information regarding the transmission history of individuals in an epidemic in order to optimize public health intervention to control the spread of disease. Information about the underlying transmission trees can be inferred from viral sequence data sampled from patients [1]. As sequencing of viral samples becomes increasingly routine, the necessity of scalability in inference tools becomes increasingly essential. PhyloPart is a tool that, given a phylogenetic tree inferred from viral sequence data, clusters the samples from which the tree was constructed such that clusters define clades in the tree and the median patristic distance between the leaves in every cluster is below a given threshold [2]. Like PhyloPart, ClusterPicker is a tool that clusters viral samples such that clusters define clades in the tree, but such that the maximum pairwise *p*-distance between sequences is below a given threshold, and ClusterPicker is able to perform the clustering significantly faster [3]. HIV-TRACE is a tool that clusters viral samples such that two individuals are placed in the same cluster if the pairwise Tamura-Nei 93 (TN93) distance between their sequences is below a given threshold [4]. As the number of sequences grows, HIV-TRACE is able to perform the clustering faster than ClusterPicker.

While these tools are able to scale to cluster thousands, or even tens of thousands, of sequences in reasonable computation time, as more patients’ viral samples are sequenced, scalability of transmission cluster inference becomes increasingly important. Here, I introduce a new tool, TreeCluster, that performs transmission cluster inference on a given phylogenetic tree, but orders of magnitude faster than existing methods.

## Implementation

TreeCluster is implemented in Python 3 and utilizes the Biopython module [5]. Given a distance threshold *t*, a branch support threshold *s*, and a bifurcating phylogenetic tree *T* with *n* leaves, TreeCluster minimizes the number of clusters of leaves of *T* in one of the following modes:

<u>Average Clade Mode:</u> The average patristic distance between leaves in the cluster is below *t*, the leaves of the cluster cannot be connected by branches with support below *s*, and the leaves of the cluster must define a clade in *T.* A post-order traversal is performed on *T* in which the average patristic distance between the leaves below each internal node *u* is computed. Then, a pre-order traversal is performed on *T*, and if the average patristic distance between leaves below a given node *u* is at most *t*, the leaves of the clade rooted at *u* is output as a cluster. This mode is O(*n*).

<u>Length:</u> The path between any two leaves in a given cluster cannot contain any edges longer than *t* or with support below *s*. A post-order traversal is performed on *T*, and if the incident edge on a given node *u* exceeds *t*, the leaves below *u* are output as a cluster. This mode is O(*n*).

<u>Length Clade:</u> Same as Length mode, but with the added restriction that clusters must define a clade in *T.* A post-order traversal is performed on *T*, and if either of the child branches of a given node *u* exceeds *t*, a cluster is output for the leaves below each of the two children of *u*. This mode is O(*n*).

<u>Maximum:</u> The maximum patristic distance between two leaves in a given cluster cannot exceed *t*, and the path between any two leaves in the cluster cannot contain edges with support below *s*. A post-order traversal is performed on *T* in which, for each node u, the largest distance from *u* to a leaf is maintained. If the longest distance to a leaf in *u*’s left subclade plus the longest distance to a leaf in *u*’s right subclade exceeds *t*, the subclade with the longer leaf distance is output as a cluster. This mode is O(*n*).

<u>Maximum Clade:</u> Same as Maximum mode, but with the added restriction that clusters must define a clade in *T.* The algorithm is identical to Maximum mode, except both subclades of the given node *u* are output as clusters instead of just the subclade containing the longer leaf distance. This mode is O(*n*).

<u>Median Clade:</u> The median patristic distance between leaves in the cluster is below *t*, the leaves of the cluster cannot be connected by branches with support below *s*, and the leaves of the cluster must define a clade in *T*. A post-order traversal is performed on *T* in which a sorted list of patristic distances is maintained for each node *u.* Then, a pre-order traversal is performed on *T,* and if the median patristic distance between leaves below a given node *u* is at most *t*, the leaves of the clade rooted at *u* is output as a cluster. This mode is O(*n*^2^ log *n*).

<u>Root Distance:</u> Cluster the leaves by cutting the tree at *t* distance below the root. Branches with support below s are simply treated as infinitely long. A pre-order traversal is performed on *T*, and if the distance between a given node *u* and the root of *T* exceeds *t*, the clade below *u* is output as a cluster. This mode is O(*n*).

<u>Single Linkage Clade:</u> The leaves of each cluster must define a clade in *T*, the leaves of the cluster cannot be connected by branches with support below s, and for each internal node *u* in the clade defined by the cluster, a leaf in the left subclade of *u* must be within *t* distance of a leaf in the right subclade of *u*. A post-order traversal is performed on *T* in which, for each node u, the shortest distance from *u* to a leaf is maintained. If the shortest distance to a leaf in *u*’s left subclade plus the shortest distance to a leaf in *u*’s right subclade exceeds *t*, both subclades of *u* are output as clusters. This mode is O(*n*).

## Results and Discussion

In order to compare the runtimes of TreeCluster with existing methods, all HIV-1 subtype B *pol* sequences (HXB2 coordinates 2,253 to 3,549) were downloaded from the Los Alamos National Laboratory (LANL) database and were subsampled into sets of *n* = 100, 200, 500, 1,000, 2,000, and 5,000 sequences (10 replicates for each value of *n*). Each replicate dataset was aligned using MAFFT [6], and phylogenetic trees were inferred under the General Time Reversible (GTR) + Γ model using FastTree 2 [7]. Polytomies were then randomly resolved with 0-length branches using the ape R package, the trees were midpoint-rooted using FastRoot [8], and Shimodaira-Hasegawa-like (SH-like) branch support values were computed using FastTree 2.

ClusterPicker, HIV-TRACE, and TreeCluster were then run on each dataset using their default parameters. Due to long execution runtime, my attempts at running PhyloPart were unsuccessful. As can be seen in Figure 1, for datasets as small as *n* = 500, TreeCluster is significantly faster than both ClusterPicker and HIV-TRACE, and as *n* increases, the gap between execution time widens by orders of magnitude.

**Figure 1.**
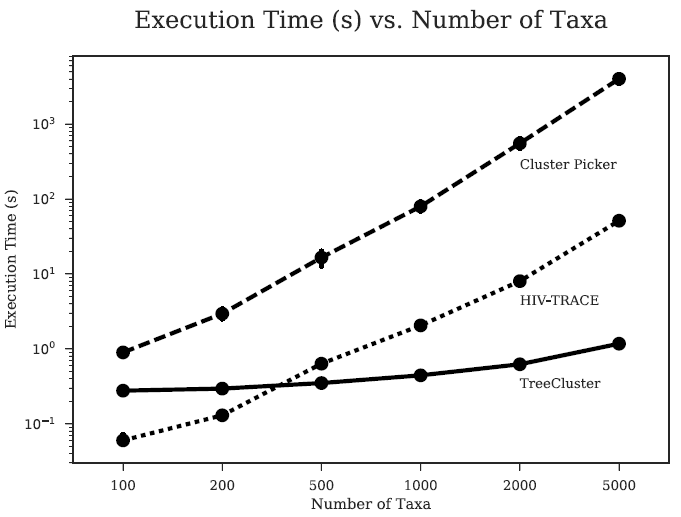
Execution times of ClusterPicker, HIV-TRACE, and TreeCluster in log-scale. Execution times (in seconds) are shown for each tool for various values of *n* sequences, with 10 replicates for each *n*. All programs were run on a CentOS 5.8 machine with an Intel Xeon X7560 2.27 GHz CPU.

## Conclusions

TreeCluster is a tool for inferring transmission clusters from a given phylogenetic tree inferred from viral sequences. It is open source and is implemented in Python 3, meaning it is cross-platform operable. Most importantly, it is orders of magnitude faster than similar tools and can thus be used on ultra-large datasets.

## Abbreviations

<u>GTR</u>: General Time Reversible
*HIV*: Human Immunodeficiency Virus
<u>LANL</u>: Los Alamos National Laboratory
<u>*pol*</u>: DNA polymerase gene
<u>SH</u>: Shimodaira-Hasegawa
<u>TN93</u>: Tamura-Nei 93

## Declarations

### Acknowledgements

I would like to acknowledge Siavash Mirarab for his guidance. I would also like to acknowledge Manon Ragonnet-Cronin and Joel Wertheim for their insights into transmission clustering.

### Availability of data and materials

TreeCluster is available at the following repository: https://github.com/niemasd/TreeCluster

The datasets used to benchmark TreeCluster are available at the following repository: https://github.com/niemasd/CP2-Paper

### Authors’ contributions

NM completed the entirety of the work.

### Consent for publication

Not applicable.

### Ethics approval and consent to participate

Not applicable.

